# Signals Generated by Neutrophil Receptors for Danger Molecules Transactivate Allosterically Modulated FFA2R: Distinct response patterns are mediated by modulators recognized by different allosteric receptor sites

**DOI:** 10.1101/2022.04.29.489985

**Authors:** Simon Lind, Kenneth L. Granberg, Huamei Forsman, Claes Dahlgren

**Author notes:** Corresponding author: Simon Lind, Department of Rheumatology and Inflammation Research, Guldhedsgatan 10A, 413 46 Göteborg, Sweden., e-mail address.

## Abstract

Positive allosteric modulators for free fatty acid receptor 2 (FFA2R/GPR43), that affect receptor function through binding to two distinct allosteric binding sites, were used to determine the correlation between the responses induced in neutrophils by two distinct activation modes; FFA2R was activated either by the orthosteric agonist propionate or by a receptor transactivation mechanism that activated FFA2R from the cytosolic side of the neutrophil plasma membrane by signals generated by the neutrophil PAFR (receptor for platelet activating factor), P2Y_2_R (receptor for ATP), FPR1 (receptor for fMLF) and FPR2 (receptor for WKYMVM). We show that the transactivation signals that activate FFA2R in the absence of any orthosteric agonist were generated downstream of the signaling G protein that couple to PAFR and P2Y_2_R. This transactivation of allosterically modulated FFA2Rs, by signals generated by PAFR/P2Y_2_R, represents a novel mechanism by which a G protein coupled receptor can be activated. Weak correlations were obtained when the FFA2R activity was induced by the transactivation signals generated by PAFRs and P2Y_2_Rs were compared with the FFA2R activity induced by the orthosteric agonist propionate. Comparison of the responses for each allosteric modulator revealed that the ratio values, calculated from the peak values of the ATP and propionate responses, varied from 0.2 to 1. Depending on the allosteric modulator, the response induced by the two different mechanisms (orthosteric activation and receptor transactivation, respectively), was equal or the propionate response was more pronounced. Importantly, we conclude that FFA2R activation from outside (orthosteric activation) and inside (receptor cross-talk/transactivation) can be selectively affected by an allosteric FFA2R modulator.

1. The allosterically modulated FFA2R is transactivated by signals generated by other GPCRs.
2. The PAF and ATP receptors transactivate FFA2R from the cytosolic side of the membrane.
3. The mechanisms that regulates activation of FFA2R from outside and inside differ.

## 1. Introduction

The short chain free fatty acid receptor 2 (FFA2R/GPR43) is a member of the large family of G protein coupled receptors (GPCRs; also known as 7-transmembrane receptors, 7TMR) expressed in neutrophils and the receptor has been suggested to have important roles in the regulation of inflammation (1, 2). The GPCRs have a common basic structure with a peptide chain spanning the membrane seven times and these receptors constitute the basis for how cells recognize specific signaling molecules in their environment (3, 4). Generally, parts of the receptor exposed on the cell surface specifically recognize agonists, while cytosolic parts of the receptor initiate/transfer the agonist induced signaling events that regulate a large number of biological processes down-stream of the activated receptor (5-7). The agonist-interaction and receptor signaling scheme was for long assumed to be some type of on/off switch, but this has during recent years been shown to be an oversimplification. This is illustrated not only by the fact that some agonists are biased and trigger the recognizing receptor to preferentially activate one signal-transduction pathway over another, but also by the fact that many (if not all) receptors have regulatory (allosteric) binding sites that are both structurally and physically separated from the orthosteric binding sites (6, 8). Such allosteric binding sites recognize non-orthosteric ligands (modulators) that change the receptor activities induced by orthosteric agonists. Propionate is an orthosteric FFA2R agonists, but the activity of the FFA2R can also be modulated by non-activating ligands binding to allosteric sites (9-11). Accordingly, allosteric FFA2R modulators transfer the natural low-activating orthosteric agonist propionate to a potent activating ligand. The original paradigm for GPCR down-stream signaling stated that a receptor specific agonist activates only one particular receptor and a receptor specific allosteric modulator affects only signaling induced by orthosteric agonists that are recognized by the modulated receptor (12). However, these limitations have been challenged and activation is now known to be triggered both by promiscuous agonists that are recognized by more than one receptor (13), as well as by receptor transactivation/cross-talk signals that activate an allosterically modulated receptor without the involvement of any orthosteric agonist that is recognized by the modulated receptor (14). In neutrophils, this type of receptor cross-talk activation/transactivation was originally shown to be a way to transfer desensitized receptors from a non-signaling to a signaling state. This process has been extensively studied in neutrophils with desensitized Formyl Peptide Receptors (FPRs) that are reactivated/transactivated by cross-talk signals generated by other pattern recognition neutrophil GPCRs (15-18). The identity of the signals that transactivate desensitized FPRs have not yet been identified, but the results obtained suggest that the transactivation signals are generated on the cytosolic side of the membrane, down-stream of the G protein coupled to the transactivating receptor partner.

We have previously shown that the positive allosteric FFA2R modulators Cmp58 and AZ1729 are recognized by two distinct different allosteric receptor sites (8, 9). These modulators transfer propionate to a potent neutrophil activating agonist. In addition, these modulators affect also the responses induced by ATP (an agonist specific for P2Y_2_R) and the two formyl peptide receptors (FPR1 and FPR2) (10, 11, 19). The mechanism for the receptor transactivation of the allosterically modulated FFA2R, by signals generated by P2Y_2_R/FPRs has notable similarities with the receptor cross-talk/transactivation of desensitized FPRs. This is illustrated by, i) the P2Y_2_R/FPR mediated responses are inhibited by FFA2R antagonists and, ii) the FFA2R activation, when mediated by the Gα_q_ coupled receptor for ATP, is abolished by an inhibitor specific for Gα_q_. These observations suggest that the allosterically modulated FFA2Rs are activated from inside the plasma membrane, by signals generated down-stream of the Gα_q_ containing G protein coupled to P2Y_2_R (10, 19).

We recently characterized several structurally diverse compounds that, based on their positive modulating effects on the neutrophil response induced by propionate, were classified as allosteric FFA2R modulators (9). These allosteric modulators interact with either of two distinctly different allosteric modulation sites in FFA2R and we herein used these as molecular tools to determine the link (if any) between activation by the orthosteric agonist propionate and by the receptor transactivation signals generated on the inside of the plasma by cross-talking GPCRs. We show that in addition to the agonist/receptor pairs (i.e., ATP/P2Y_2_R, fMLF/FPR1 and WKYMVM/FPR2) earlier shown to transactivate FFA2R, also the receptor for platelet activating factor (PAFR) generates signals that transactivate FFA2R. When comparing transactivation of FFA2R mediated by the Gα_i_ coupled FPR1 and FPR2, the activation patterns were very similar to those mediated by the P2Y_2_R, FPR1 and FPR2. Although not as conclusive, there were also similarities between how the allosteric modulators affect the Gα_q_ coupled PAF and ATP induced neutrophil NADPH oxidase activity. More importantly, the positive allosteric effect was selective for propionate suggesting that the mechanisms that regulate activation of the receptor from outside (orthosteric activation) and inside (cross-talk/transactivation) are different.

## 2. Material and Methods

### 2.1. Chemicals

Isoluminol, TNF-α, fMLF, ATP, PAF and propionic acid, were purchased from Sigma (Sigma Chemical Co., St. Louis, MO, USA). Dextran and Ficoll-Paque were obtained from GE-Healthcare Bio-Science (Uppsala, Sweden). Horseradish peroxidase (HRP) was obtained from Boehringer Mannheim (Mannheim, Germany). The PAF agonist were from Avanti Polar Lipids Inc. (Alabama, USA), and all transactivating receptor agonist tools are described in Table 1. The allosteric FFA2R modulator Cmp58 ((*S*)-2-(4-chlorophenyl)-3,3-dimethyl-*N*-(5-phenylthiazol-2-yl)butanamide, the FFA2R antagonists GLPG0974 (4-[[(*R*)-1-(benzo[*b*]-thiophene-3-carbonyl)-2-methyl-azetidine-2-carbonyl]-(3-chlorobenzyl)amino]-butyric acid) and AR-C118925 {5-[[5-(2,8-dimethyl-5*H*-dibenzo[*a,d*]cyclohepten-5-yl)-3,4-dihydro-2-oxo-4-thioxo-1(*2H*)-pyrimidinyl]methyl]-*N*-*2H*-tetrazol-5-yl-2-furancarboxamide} were obtained from Calbiochem-Merck Millipore (Billerica, USA) and TOCRIS (Bristol, UK). The antagonist CATPB ((*S*)-3-(2-(3-chlorophenyl)acetamido)-4-(4-(trifluoromethyl)phenyl) butanoic acid was synthesized as described previously (20, 21) and obtained (as generous gifts) from Trond Ulven (Odense University, Denmark). The Gα_q_ inhibitor YM-254890 was purchased from Wako Chemicals (Neuss, Germany). The FPR2 specific hexapeptide Trp-Tyr-Met-Val-Met-NH2 (WKYMVM) was synthesized and purified by HPLC by Alta Bioscience (University of Birmingham, Birmingham, United Kingdom). The allosteric FFA2R modulator AZ1729 (9, 11, 22) together with all the other FFA2R ligands included in the study (Fig 2) were provided by AstraZeneca (Gothenburg, Sweden).

**Table I.** Classification of FFA2R ligands in neutrophils

| Compound | Activation<sup>†</sup> | Allosteric modulation of |  |  | Classification |
| --- | --- | --- | --- | --- | --- |
|  |  | Propionate | AZ1729 | Cmp58 |  |
| Cmp58 | - | + | + | - | Cmp58 AM* |
| AZ5994 | + | + | + | - | Cmp58-like agonist |
| AZ7004 | - | + | + | - | Cmp58-like AM |
| CATPB | - | - | - | - | Antagonist |
| GLPG0974 | - | - | - | - | Antagonist |
| AZ1729 | - | + | - | + | AZ1729 AM |
| AZ0688 | - | + | - | + | AZ1729-like AM |
| AZ0682 | - | + | - | + | AZ1729-like AM |
| AZ8703 | - | + | - | + | AZ1729-like AM |
| AZ1702 | - | + | - | + | AZ1729-like AM |

† Indicates agonistic properties in neutrophils \*AM: Allosteric modulator

Subsequent dilutions of receptor ligand and other reagents were made in Krebs-Ringer Glucose phosphate buffer (KRG; 120 mM NaCl, 4.9 mM KCl, 1.7 mM KH_2_PO_4_, 8.3 mM NaH_2_PO_4_, 1.2 mM MgSO_4_, 10 mM glucose, and 1 mM CaCl_2_ in dH_2_O, pH 7.3).

### 2.2. Isolation of human neutrophils

Neutrophils were isolated from buffy coats from healthy blood donors by dextran sedimentation and Ficoll-Paque gradient centrifugation as described by Bøyum (23). Remaining erythrocytes were removed by hypotonic lysis and the neutrophils were washed and resuspended in KRG. More than 90% of the cells were neutrophils with a viability of >95%. To amplify the activation signals, the neutrophils were primed with TNF-α (10 ng/mL for 20 min at 37°C), and then stored on ice until use.

### 2.3. Measuring NADPH-oxidase activity

Isoluminol-enhanced chemiluminescence (CL) technique was used to measure superoxide production (24), the precursor of production of reactive oxygen species (ROS), by the NADPH-oxidase activity as described (25, 26). In short, the measurements were performed in a six-channel Biolumat LB 9505 (Berthold Co., Wildbad, Germany), using disposable 4 mL polypropylene tubes and a 900 μL reaction mixture containing 10^5^ neutrophils, isoluminol (0.2 μM) and HRP (4 Units/mL). The tubes were equilibrated for 5 min at 37°C, before addition of agonist (100 μL) and the light emission was recorded continuously over time. In experiments where the effects of receptor specific antagonists were determined, these were added to the reaction mixture 1–5 min before stimulation and control neutrophils were separately incubated under the same conditions but in the absence of an antagonist.

### 2.4. Statistical analysis

Statistical calculations were performed in GraphPad Prism 9.3 (Graphpad Software, San Diego, CA, USA). Which specific statistical tests that are performed are stated in the relevant figure legend. A *p*-value < 0.05 was regarded as statistically significant difference and is indicated by *p < 0.05, \*\**p* < 0.01, \*\*\**p* < 0.001. Correlation between the different modes of activation was performed and presented with Pearson coefficients and plotted together with simple linear regression showing 95% confidence intervals. To further study the agreement a Bland Altman plot (PMID: 2868172) was calculated showing bias and the limits of the agreement. Some data was also further statistically analyzed using a one-way ANOVA followed by Dunnett’s multiple comparison or, paired Student’s *t*-test. Even though the results in the figures are presented as percent of control values, the statistics have been calculated using the raw data.

### 2.5. Ethics Statement

In this study, conducted at the Sahlgrenska Academy in Sweden, buffy coats obtained from the blood bank at Sahlgrenska University Hospital, Gothenburg, Sweden have been used. According to the Swedish legislation section code 4§ 3p SFS 2003:460 (Lag om etikprövning av forskning som avser människor), no ethical approval was needed since the buffy coats were provided anonymously and could not be traced back to a specific donor.

## Results

### 4.1 Two distinct different mechanisms used by FFA2R to activate the neutrophil NADPH-oxidase

#### 4.1.1. Activation by the orthosteric FFA2R agonist propionate (Fig 1)

In order for propionate, an orthosteric FFA2R agonist, to activate the neutrophil electron transporting NADPH-oxidase system that produces superoxide anions (O_2_^-^), the receptor must be allosterically modulated (see (9) and (27)). In agreement with the receptor-selectivity of propionate, the response induced by this agonist in the presence of either of the two allosteric modulators Cmp58 and AZ1729 (Fig 2), was inhibited by an FFA2R antagonist; for comparison the response was not affected by an antagonist specific for P2Y_2_R, the neutrophil receptor for ATP (Fig 3A and B). The allosteric modulators Cmp58 and AZ1729 have been shown to be specific for FFA2R (22, 28) and to be recognized by two different allosteric receptor sites (9, 11). The response induced by propionate was not affected by the Gα_q_ inhibitor YM-254890 (Fig 3B), showing that no Gα_q_ containing G protein is involved in the down-stream signaling of propionate activated FFA2Rs.

**Figure 1.**
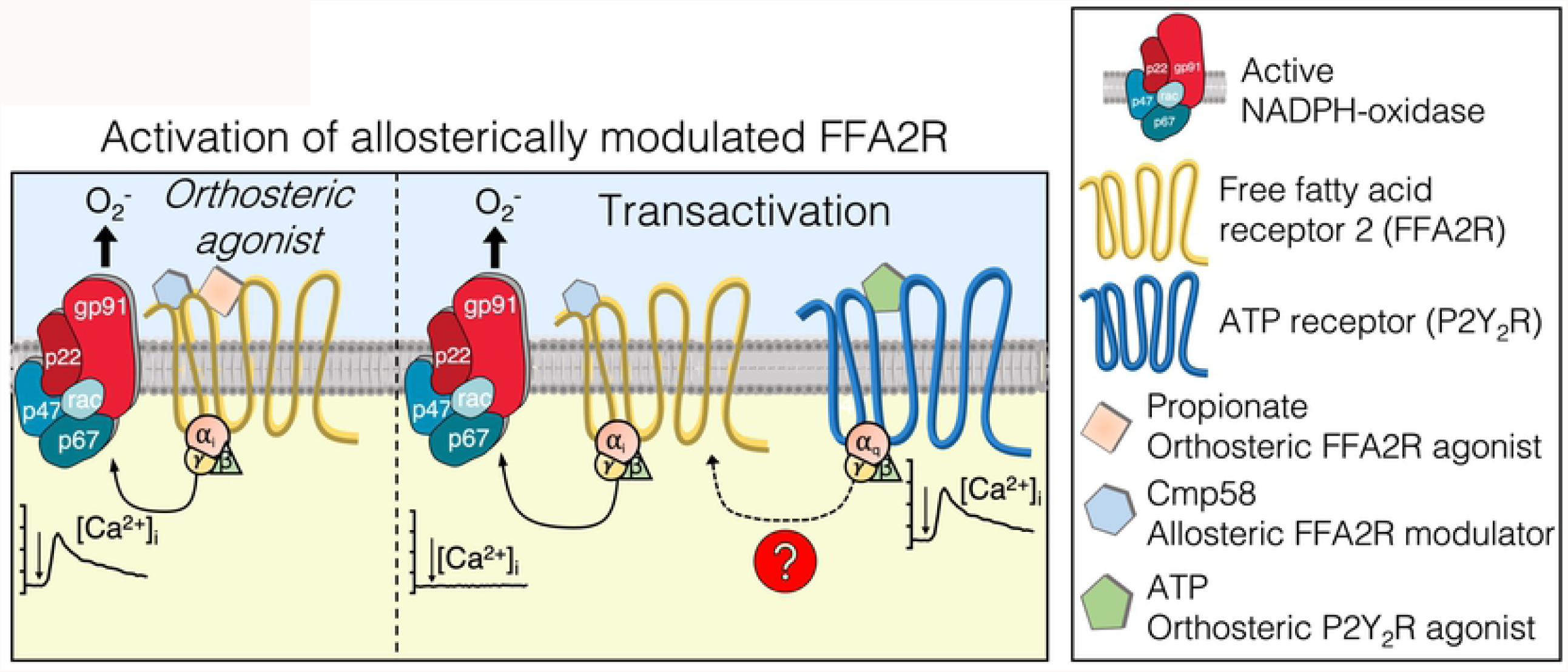
Activation of allosterically modulated FFA2R by two different mechanisms. ***Left***: Activation of FFA2R by the orthosteric agonist propionate includes a transient rise in [Ca^2+^]_i_ and production of O_2_^-^ (left). **Right**: Transactivation of FFA2R by signals generated by the ATP receptor (inducing a rise in [Ca^2+^]_i_ by itself) leads to production of O_2_^-^ but no FFA2R induced transient rise in [Ca^2+^]_i_.

**Figure 2.**
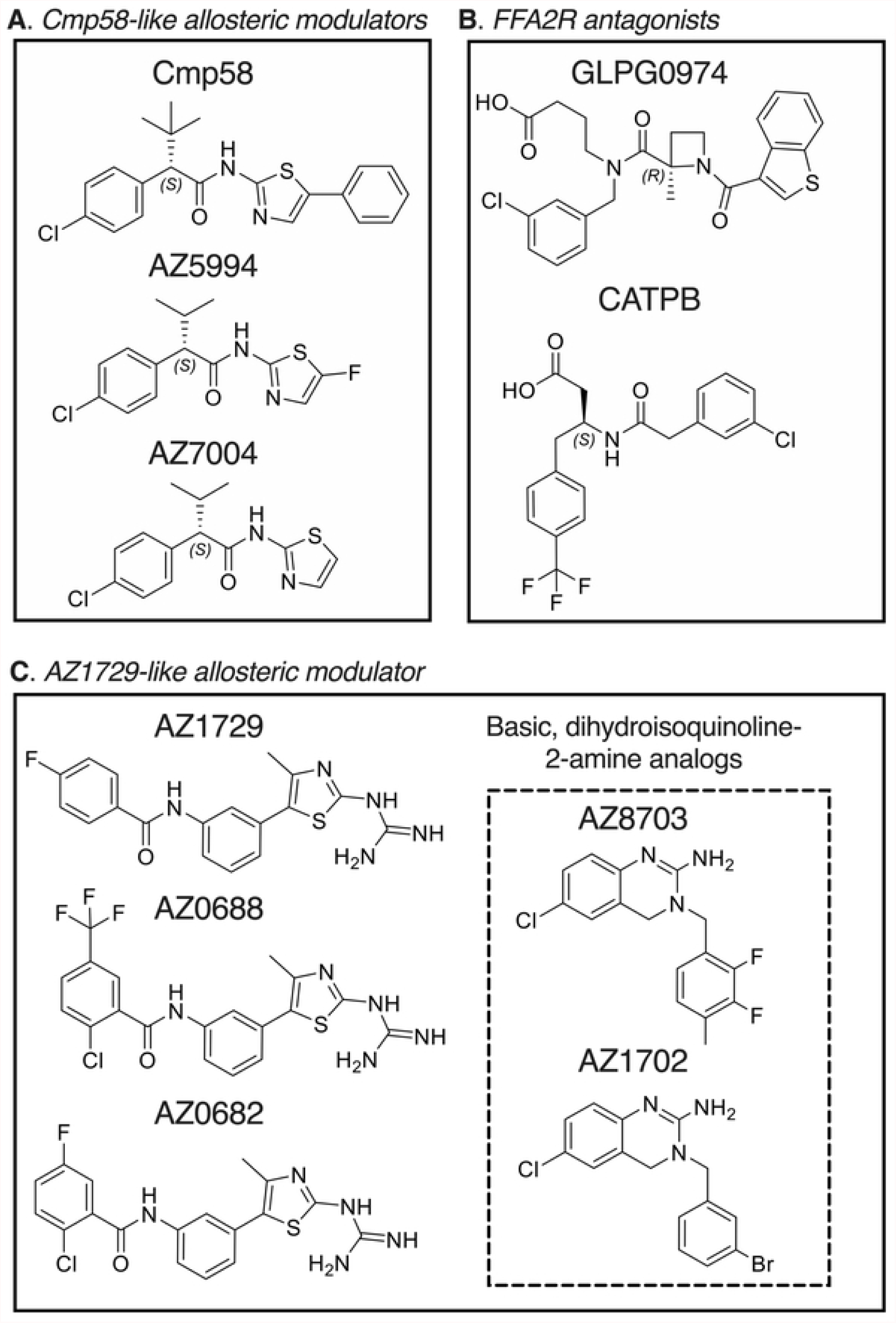
Chemical structures of the FFA2R ligands used in the study. Structural variants of allosteric FFA2R modulators with distinct interaction properties were chosen through screening of a mini-library of small compounds. Ten compounds were identified as allosteric FFA2R modulators. Two selective FFA2R antagonists, GPLG0974 and CATPB, were also selected.

**Figure 3.**
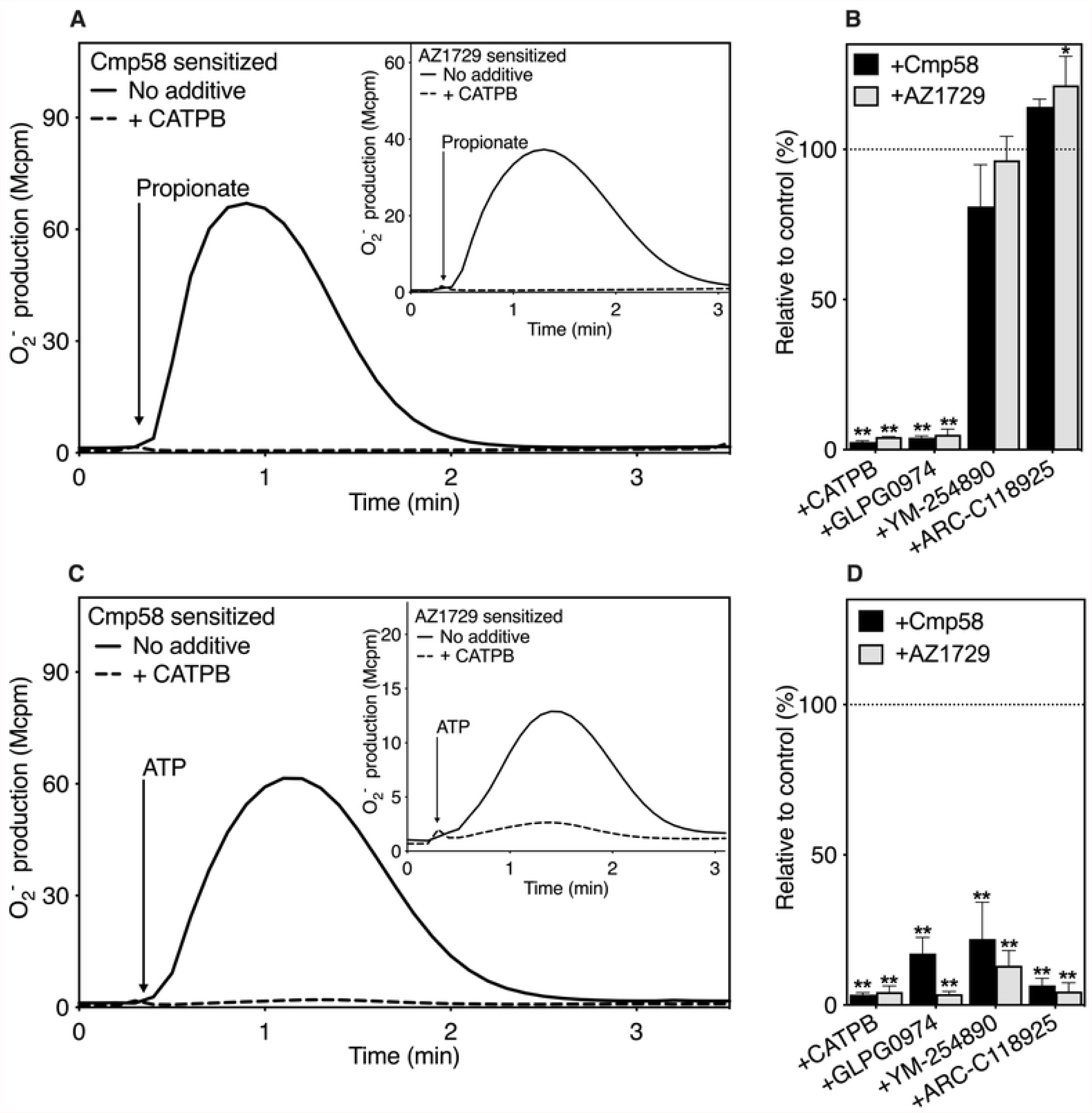
Selective inhibition by specific antagonists/inhibitors on the neutrophil FFA2R dependent response when triggered by the orthosteric FFA2R agonist propionate and the P2Y2R agonist ATP, respectively. (**A**) Neutrophils sensitized with Cmp58 (1 μM for 5 min) were activated (indicated with arrow) with the FFA2R orthosteric agonist, propionate (25 μM solid line). The release of O_2_^-^ was measured continuously and expressed in Mcpm. The response induced by propionate in Cmp58 sensitized (1 μM for 5 min) neutrophils was also treated at the same time with the FFA2R specific antagonists, CATPB (dashed line, 100 nM for 5 min). One of four representative experiment is shown. Inset: Same experimental setup was used but neutrophils were instead sensitized to FFA2R specific allosteric modulator, AZ1729 (1 μM for 5 min). (**B**) The peak O_2_^-^ production values were determined and the ratios between the propionate modulated responses from Cmp58 (1 μM) or AZ1729 (1 μM) in the absence and presence of one the FFA2R antagonist CATPB (100 nM) and GLPG0974 (100 nM), the Gα_q_ inhibitor YM-254890 (200 nM), the selective P2Y_2_R antagonist AR-C118925 (1 μM) were calculated and expressed in remaining activity (in percent) in the presence of the respective antagonist (mean + SD, n = 4). (**C, inset**) Same experimental setup as in (**A**) but the activating ligand is instead the P2Y_2_ agonist, ATP (10 μM). (**D**) The peak O_2_^-^ production values were determined and the ratios between the responses in the ATP modulated responses from Cmp58 (1 μM) or AZ1729 (1 μM) in absence and presence of the respective antagonist CATPB (100 nM) or GLPG0974 (100 nM), YM-254890 (200 nM), and AR-C118925 (1 μM) were calculated and expressed in remaining activity (in percent) in the presence of the respective antagonist (mean+SD, n = 4). Statistical analyses were performed (**B** and **D**) using a one-way ANOVA followed by a Dunnett’s multiple comparison test comparing the peak responses in the absence and presence of respective inhibitor.

#### 4.1.2. Activation by ATP, an orthosteric P2Y_2_R agonist (Fig 1)

No activation of the O_2_^-^ producing NADPH-oxidase was induced by the P2Y_2_R agonist ATP in naive neutrophils (earlier shown in (10, 11, 19)). In the presence of either of the allosteric FFA2R modulators Cmp58 or AZ1729, ATP is a potent activating agonist (Fig 3C). The differences in the inhibition patterns of a selection of inhibitors/antagonists show that the orthosteric FFA2R agonist propionate and the P2Y_2_R agonist ATP trigger different pathways to activate FFA2R. The ATP induced response was in contrast to the propionate induced response, inhibited both by the P2Y_2_R specific antagonist AR-C118925 and by the Gα_q_ selective inhibitor YM-254890 (Fig. 3D). In addition, and irrespectively of the allosteric FFA2R modulator included, also the ATP induced response was fully inhibited by the FFA2R specific antagonists CATPB and GLPG0974 (Fig 3C and D).

Taken together, these data show that an activation of allosterically modulated FFA2Rs can be achieved by two different mechanisms; i) the NADPH-oxidase activating signals, induced by a conventional FFA2R agonist recognized by the orthosteric receptor binding pocket exposed on the cell surface, were generated independent of any Gα_q_ containing G protein, and ii) the FFA2R dependent activation of the NADPH-oxidase, mediated by signals generated by P2Y_2_R, was achieved without the involvement of any orthosteric FFA2R agonist and were generated down-stream of a Gα_q_ containing G protein.

### 4.2. Ligand tools for FFA2R

By definition, allosteric FFA2R modulators should positively modulate the activity induced by an orthosteric FFA2R agonist such as propionate. We have earlier descibed eight compounds including Cmp58 and AZ1729, that all fulfill the basic criteria for an allosteric FFA2R modulator ((9); Fig 2; Table 2); i.e., they lack direct neutrophil activating effects alone but turn propionate into a neutrophil activating agonist. Based on an orthosteric independent activation mechanism induced by the non-activating modulators when combined with either Cmp58 or AZ1729 (11), we have shown that FFA2R has two different allosteric binding site and the modulaters used in this study were classified as “Cmp58 like” (activation achieved only together with AZ1729) or as “AZ1729 like” (activation achieved only together with Cmp58) (Table 2). All the allosteric FFA2R modulators (in total eight; Fig 2) were used as tools to determine differences (if any) related to the activation mechanism by which the allosteric modulated FFA2Rs were activated by, i) a direct binding of the orthosteric FFA2R agonist propionate that initiates Gα_q_ independent activation signals, and ii) by the receptor cross-talk/transactivation mechanism shown to be initiated for example by binding of ATP to P2Y_2_R that generates activting signals downstrem of a Gα_q_ containg G protein.

**Table II.** Classification of orthosteric agonists and their respective GPCR

| Agonist | Recognizing receptor (GPCR) |
| --- | --- |
| Propionate | Free Fatty Acid receptor 2 (FFA2R) |
| ATP | P2Y purinoceptor 2 (P2Y<sub>2</sub>R) |
| PAF | Platelet Activating Factor Receptor (PAFR) |
| fMLF | Formyl Peptide Receptor 1 (FPR1) |
| WKYMVM | Formyl Peptide Receptor 2 (FPR2) |

### 4.3. Relation between the neutrophil responses when FFA2R was activated by two different activation mechanisms

#### 4.3.1. Correlation between the FFA2R transduced neutrophil responses induced by propionate and ATP

Previous studies (10, 19) and Fig 3 show that the specific P2Y_2_R agonist ATP as well as the specific FFA2R agonist propionate activate FFA2R provided that the receptor is allosterically modulated. However, the activation mechanisms differ - whereas propionate is an orthosteric FFA2R agonist that transfers FFA2R to a signaling state by binding to the orthosteric FFA2R binding site exposed on the cell surface, the FFA2R activation mechanism initiated by ATP binding to its neutrophil receptor, rely on signals generated down-steam of the G protein coupled to P2Y_2_R. The eight allosteric FFA2R modulators (Table 2) were used to determine the direct relation between the neutrophil responses when FFA2R was activated by the “propionate and ATP mechanism”, respectively. The allosteric FFA2R modulating compounds were allowed to sensitize neutrophils during a 5 min incubation with respective compound. The neutrophils were then activated with propionate (25 μM) or ATP (10 μM) and the production of superoxide (O_2_^-^) was followed over time. The peak values of the responses were determined, and eight independent experiments were performed.

There are multiple ways to determine association between two responses. A direct application of the correlation coefficient (*r*) was used to determine the direction (positive or negative) and quantify the strength of the association (if any) between the propionate and ATP induced responses. Accordingly, to determine the presumable linear correlation between responses of allosterically modulated FFA2Rs when triggered by propionate and ATP, respectively, a scatter plot was generated followed by a linear regression curve as well as a Pearson correlation calculation of r and R^2^) was performed. The peak response values from all experiments performed with the different allosteric modulators were included in the plot (Fig 4A). The linear regression and the Pearson correlation value (r≈0.72), supports a moderate to strong positive linear correlation between the two data sets. A portion of data points were, however, not in the area that defined the 95% confidence interval (marked by red lines in Fig 4A). The R^2^ ≈ 0.52 means that approximately half of the observed variation could be explained by the model.

**Figure 4.**
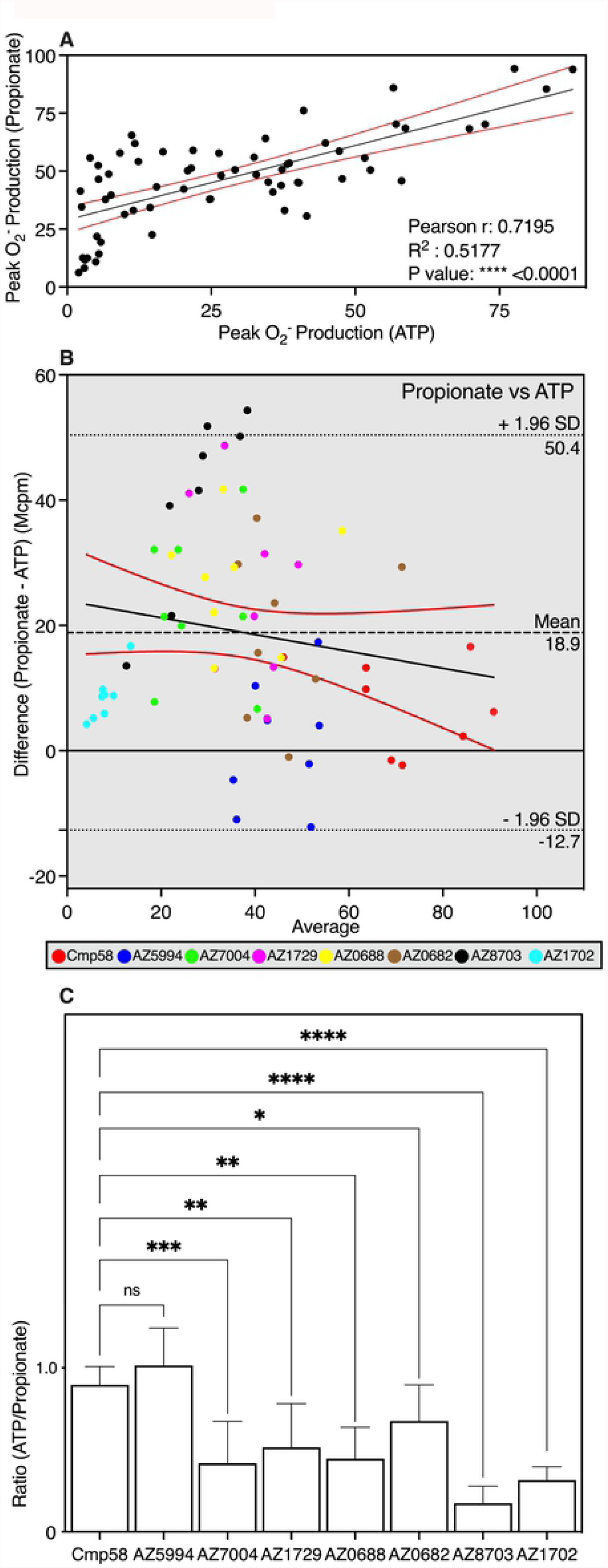
Neutrophil activation by propionate or ATP in neutrophils sensitized with allosteric FFA2R modulators. (**A**) Scatter plot for correlation of peak Mcpm value between propionate (25 μM) and ATP (10 μM) of all FFA2R modulator responses collected. Black solid line corresponds to a linear regression fit red solid lines correspond to 95% confidence slope for the linear regression fit. Pearson coefficients are also given. (**B**) Bland-Altman analysis of the peak Mcpm values between propionate and ATP for all FFA2R modulator responses collected that were used to estimate the agreement between the readouts. The Bland-Altman plot for the differences (propionate – ATP) in Mcpm values between the two responses against the averages (peak values of (propionate + ATP)/2). Line of equality is presented as black dashed line. The 95% upper and lower CI of limits of agreement is plotted as black dotted line. Linear regression line of differences is drawn as black line with red lines show the 95% confidence interval. Each allosteric modulators response for propionate and ATP is color-coded to a unique color in the Bland-Altman plot. (**C**) Human neutrophils sensitized with allosteric FFA2R modulators (1 μM each) were activated by propionate (25 μM) or ATP (10 μM) and O_2_^-^ production was measured continuously. The peak activities were determined of the propionate and ATP induced responses in neutrophils for each allosteric modulator and then presented as a ratio of the two responses. One-way ANOVA followed by Dunnett’s post-hoc test was used to calculate significant difference deviation from Cmp58. Eight independent experiments with cells isolated from healthy donors were performed and the results are given as mean + SD.

#### 4.3.2. Agreement between the FFA2R transduced responses induced by propionate and ATP analyzed by the Bland-Altman technique

The correct statistical method to evaluate the level of agreement between two data sets is not evident. Commonly, a correlation coefficient (*r*) between the results of two measurement methods has been specified as a level of agreement. An alternative approach to the conventional method was introduced by Altman and Bland in 1986 (29, 30), and this Bland-Altman plot/analysis has become frequently used as a suitable analytical tool to determining the limits of agreement between data sets obtained with two different quantitative methods (29, 30).

Also, the Bland-Altman estimation of agreement is based on a scatter data plot in which the values of the differences between paired measurements (A-B = Y value) are plotted against the mean values of the two [(A+B)/2 = X value]. Accordingly, using this equation in the Bland-Altman graph describing the effects of the different allosteric modulators, the Y values in the plot equals the propionate induced neutrophil response (peak value of O_2_^-^ production (A)) from which the corresponding ATP induced response (B) was substracted, and the X values equals the mean value of the propionate response and the corresponding ATP response (Fig 4B). This graph provides two main pieces of information; i) the average of all the differences that was 18.9 units and ii) the 95% limits of agreement with a lower limit of -12.7 and an upper of 50.4 units. A full support for the significant positive correlation suggested by the Pearson r values (Fig 4A) would require that the data points in the Bland-Altman plot lie perfectly along the line of equality. The data show that compared to ATP, the propionate response measures on the avarage an 18.9 Mcpm higher value. It is recommended that 95% of the data points should lie within ±1.96 SD of the mean difference – limits of agreement, and only two data points exceeds 50.4 Mcpm, which indicates there are agreement between the tests. Applying a simple linear correlation with confidence interval limits plot to look for proportional differences (31, 32) show that there is a slight negative bias, and taken together the data indicate that the allosteric modulators that give mid to high responses are more prone to larger differences between propionate and ATP. Even though most values fall within the limits of agreement, the different allosteric modulators clustered in different parts of the Bland-Altman plot (Fig 4B). This suggests that although there is an over all positive correlation there is not a direct agreement between the two responses for different modulators.

#### 4.3.3. The allosteric FFA2R modulators affect the propionate and ATP responses differently

A simpler and more direct way to determine the similarities/differences between the effects of the allosteric FFA2R modulators on the NADPH-oxidase activity induced by propionate and ATP, is to compare the two responses separately for each allosteric modulator (shown in Fig 4C). The ratio between the peak values of the ATP and propionate responses was close to one with Cmp58 as well as AZ5994 whereas the values for the rest of the allosteric FFA2R modulators were lower and significantly different when compared to Cmp58 (Fig 4C). Taken together, the data presented suggest that the allosteric modulators affect the propionate and ATP induced responses differently.

#### 4.3.4. Relation between the neutrophil responses when FFA2R was activated through receptor cross-talk by two Gα_q_ coupled GPCRs

##### The allosterically modulated FFA2R is transactivated by the receptor for platelet activating factor (PAFR)

Binding of the lipid neutrophil chemoattractant PAF to its neutrophil receptor triggers an activation of the neutrophil NADPH-oxidase, and similar to P2Y_2_R the receptor down-stream signaling by the PAFR relies on a Gα_q_ containing G protein (Fig 5A). Depending on the concentration of PAF, Cmp58 was without effect (100 nM concentration of PAF; Fig 5A) or turned a non-activating concentration (1 nM; Fig 5B) into a potent transactivating ligand. The PAF induced transactivation was inhibited both by the PAFR specific antagonist WEB and by the Gα_q_ selective inhibitor YM-254890. In addition, and irrespectively of the allosteric FFA2R modulator included, also the PAF induced response was inhibited by the FFA2R specific antagonists CATPB. Taken together, these data show that PAF induced activation of allosterically modulated FFA2Rs was achieved by a receptor cross-talk/transactivation mechanism, similar to that induced by ATP, without the involvement of any orthosteric FFA2R agonist and by signals generated down-stream of a Gα_q_ containing G protein coupled to the PAF receptor.

**Figure 5.**
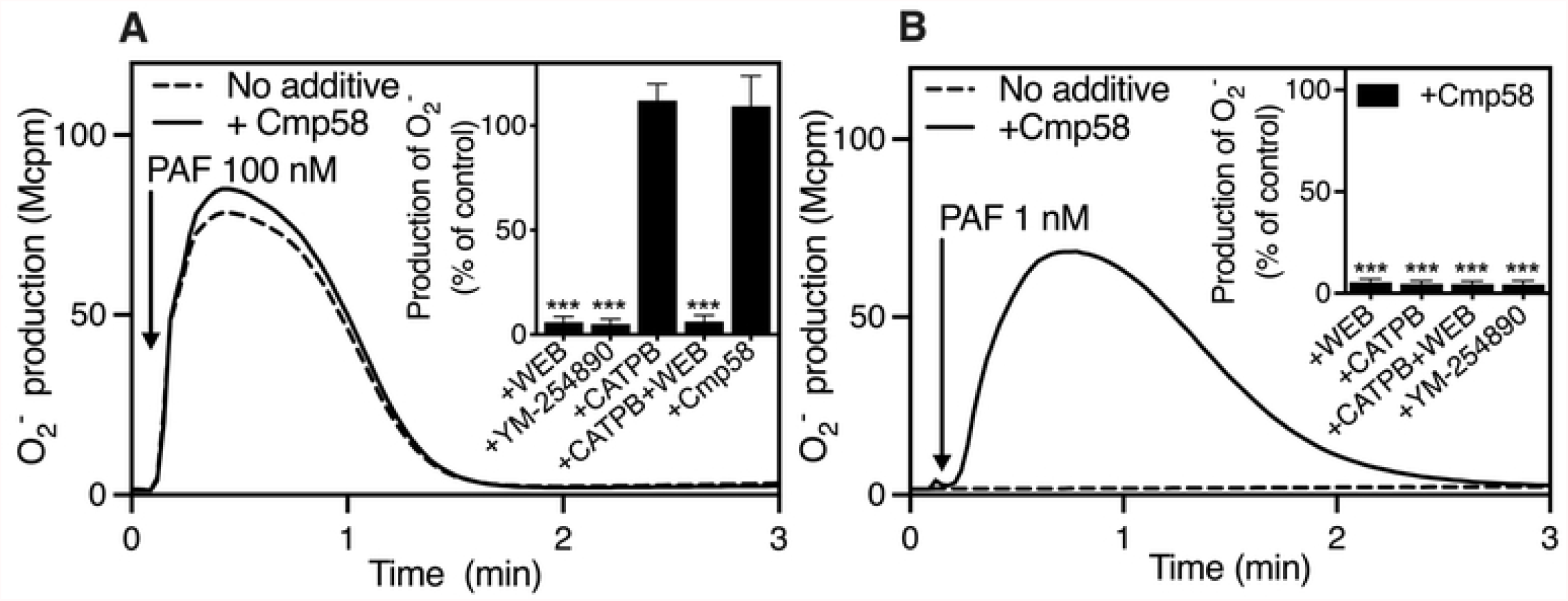
Selective inhibition by specific antagonists/inhibitors on the neutrophil FFA2R dependent cross-talk response when triggered by the orthosteric PAFR agonist PAF. (**A**) Neutrophils sensitized with Cmp58 (1 μM for 5 min solid line) or without (dashed line) were activated (indicated with arrow) with the PAFR orthosteric agonist, PAF (100 nM). The release of O_2_^-^ was measured continuously and expressed as Mcpm. One representative experiment out of three is shown. (**A**, inset) The PAF response was also treated at the same time by the PAFR specific antagonist WEB (1 μM), Gα_q_ specific inhibitor YM-254890 (200 nM), FFA2R specific antagonists, CATPB (100 nM for 5 min) as well as the combination of CAPTB (100 nM) and WEB (1 μM). The peak O_2_^-^ production values were determined and the ratios between the PAF responses in the absence and presence of the respective were calculated and expressed in remaining activity (in percent) in the presence of the respective antagonist (mean + SD, n = 3). (**B**) An experimental setup as that in **A** was used but with a 100-fold lower concentration of PAF (1 nM). (**B, inset**) Similar to (**A, inset**), however the peak O_2_^-^ production values were determined and the ratios between the responses in the PAF (1 nM) modulated responses from Cmp58 (1 μM in absence and presence of the respective antagonist/inhibitor. Statistical analyses were performed (**A** and **B, insets**) using a one-way ANOVA followed by a Dunnett’s multiple comparison test comparing the peak PAF responses in the absence and presence of respective treatment.

##### Correlation between the FFA2R transduced neutrophil responses induced by ATP and PAF

The correlation (*r*) was used to determine the direction (positive or negative) and quantify the strength of the association also between the ATP and PAF induced responses. The results obtained, when combining responses from the allosteric modulators in a scatterplot with a fitted linear regression curve as well as calulation of the Pearson coeficcients of the ATP and PAF responses (Fig 6A), revealed not only a strong positive correlation (r=0.89) but also the linear correlation was strong (R^2^ 0.79; Fig 6A). The conclusions that could be drawn from the Bland-Altman graphs, describing the effects of the different allosteric modulators on the agreement between the PAF and ATP induced responses (Fig 6B), were that i) that PAF response was almost always higher than the ATP response and ii) that there was a positive trend. All points fall in the 95% limits of agreement.

**Figure 6.**
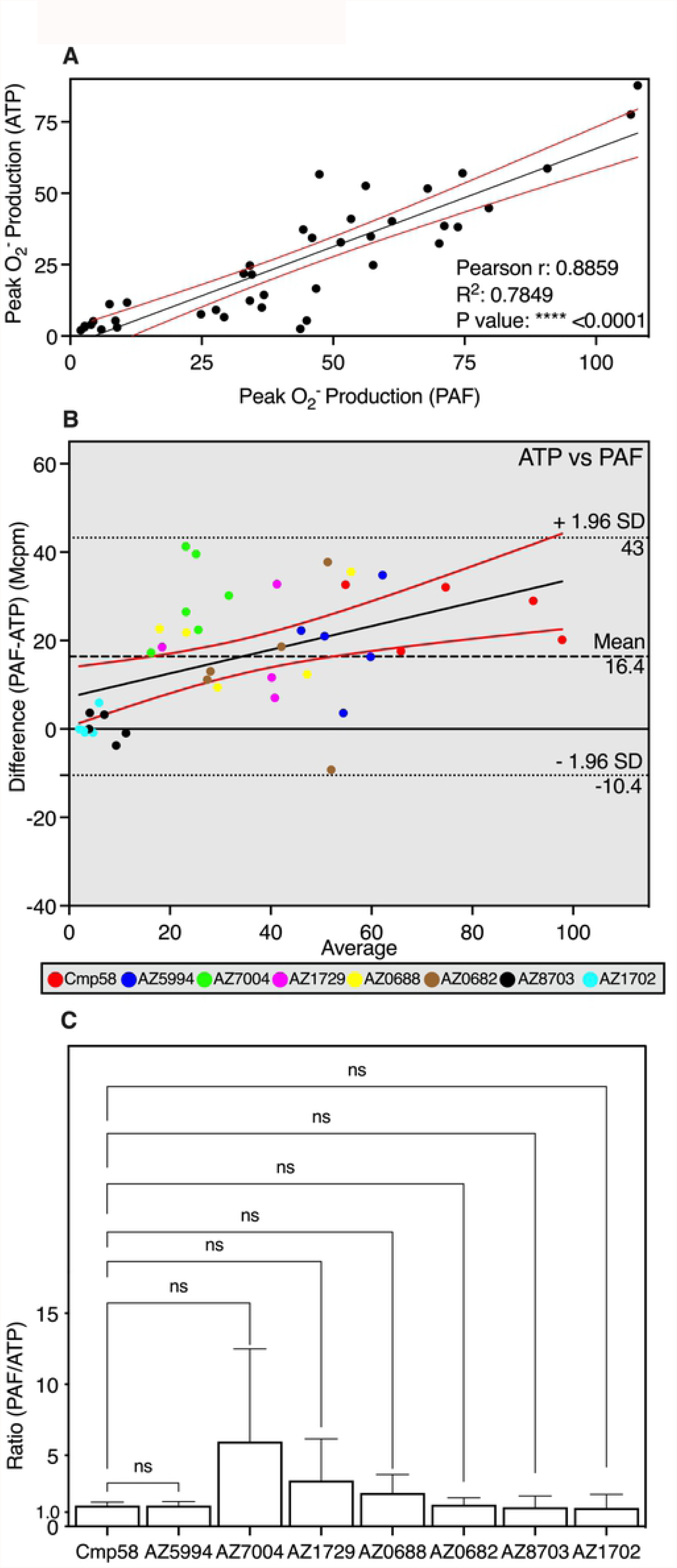
Comparison of neutrophil activation by propionate, PAF or ATP in neutrophils sensitized with allosteric FFA2R modulators. (**A**) Human neutrophils sensitized with the different allosteric FFA2R modulators (1 μM each) were activated by ATP (10 μM) or PAF (1 nM) and O_2_^-^ production was measured continuously. The peak activities were determined of the propionate and ATP induced responses in neutrophils for each allosteric modulator. Scatter plot of peak Mcpm value between ATP and PAF (**A**) of all FFA2R modulator responses collected. Black solid line corresponds to linear regression and red solid lines corresponds to 95% confidence slope for the linear regression fit. The Pearson r and R^2^ value are also given. (**B**) Bland-Altman analysis for peak Mcpm value between PAF and ATP of all FFA2R modulator responses collected was used to estimate the agreement. The Bland-Altman plots the PAF-ATP responses in Mcpm values against the averages (i.e. peak values of (PAF+ATP)/2). Line of equality is presented as black dashed line. The 95% upper and lower CI of limits of agreement is plotted as black dotted lines. The regression line of differences is drawn as a black line with red lines showing the 95% confidence interval. Each allosteric modulators response is color-coded to a unique color. (**C**) The peak O_2_^-^ activities were determined and is projected in a bar graph as the ratio between PAF (1 nM) and ATP (10 μM) for all allosteric modulators tested. Six independent experiments with cells isolated from healthy donors were performed and the results are given as mean + SD. One-way ANOVA followed by Dunnett’s post-hoc test was used to calculate significant difference deviation from Cmp58.

##### Direct correlation between the FFA2R transduced neutrophil responses

Comparing the responses for each allosteric modulator, by the more direct way to determine the similarities/differences between the effects of the allosteric FFA2R modulators on the NADPH-oxidase activity, showed when comparing the PAF and ATP induced neutrophil response responses by the more direct way, that there were no differences in the ratio between the peak values of the PAF and ATP responses (Fig 6C).

Taken together, the data suggest that even though both PAF and ATP activate the allosterically modulated FFA2R through receptor cross-talk/transactivation signals generated down-stream of the Gα_q_ coupled PAFR and P2Y_2_R, respectively, the allosteric modulated FFA2R was activated in a similar but not identical way by PAF and ATP.

#### 4.3.5. Relation between the neutrophil responses when FFA2R was activated through receptor cross-talk by two Gα_i_ coupled GPCRs

The neutrophil members of the formyl peptide receptor family (FPR1 and FPR2) sense N-formylated peptides of bacterial and mitochondrial origin (33, 34). We have earlier shown that low (normally non-activating) concentrations of FPR agonists (fMLF for FPR1 and WKYMVM for FPR2 specific) activate the O_2_^-^ producing NADPH-oxidase in the presence of an allosteric FFA2R modulator (10, 19). This activation is inhibited by FFA2R specific antagonists, suggesting that a similar type of receptor cross talk/transactivation, as that described above for P2Y_2_R/FFA2R, is the mechanism by which FPRs activate FFA2R. In contrast to the ATP and PAF induced activation, the Gα_q_ inhibitor (YM-254890) has no effect on FPR signaling. This is in agreement with the generally accepted opinion that the FPRs couple to a Gα_i_ containing G protein (35). Neutrophils sensitized for five minutes with the allosteric FFA2R modulators were activated by low (non-activating) concentrations of fMLF and WKYMVM. The data obtained from the Pearson scatterplot, comparing peak responses induced fMLF and WKYMVM through the FFA2R cross-talk mechanism, the positive linear correlation had an R^2^ value of 0.97 (Fig 7A). The agreement between fMLF and WKYMVM was obvious also when plotting the Bland-Altman scatter plots of the two peptides (Fig 7B). The bias for the agreement between WKYMVM and fMLF is 3.9 Mcpm, which consider as an acceptable level of differentiation (36) between two different modes of activation. However, the responses induced by fMLF and WKYMVM were low for the majority of the allosteric modulators (Fig 7A).

**Figure 7.**
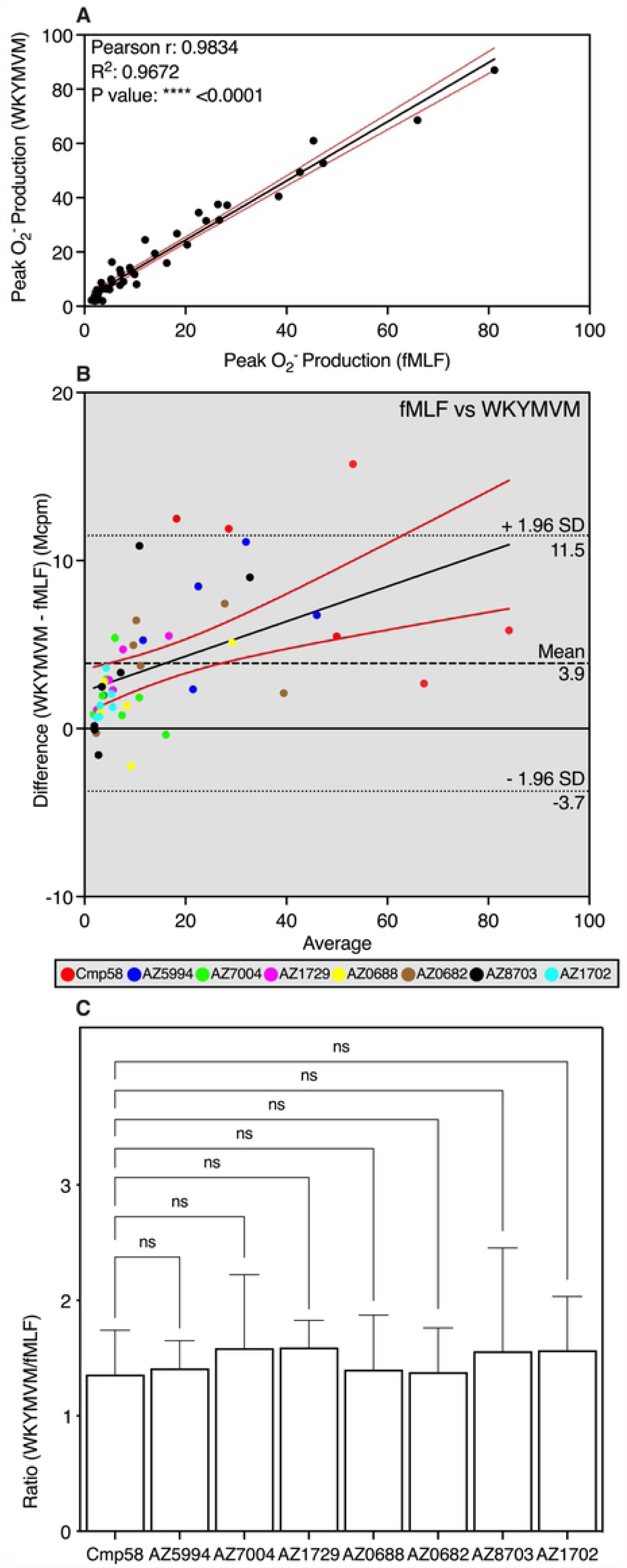
Comparison of neutrophil activation by propionate, fMLF or WKYMVM in neutrophils sensitized with allosteric FFA2R modulators. (**A/B/C**) Human neutrophils sensitized with allosteric FFA2R modulators (1 μM each) were activated by fMLF (0.5 nM) or WKYMVM (2.5 nM) and the O_2_^-^ production was measured continuously. The peak activities were determined of the fMLF and WKYMVM induced responses in neutrophils for each allosteric modulator. Scatter plot for correlation of peak Mcpm value between or fMLF and WKYMVM (**A**) of all FFA2R modulator responses collected. The black solid line corresponds to linear regression fit and red solid lines corresponds 95% confidence slope for the linear regression fit. r and R^2^ value are also given. (**B**) Bland-Altman analysis for peak Mcpm value between fMLF and WKYMVM of all FFA2R modulator responses collected was used to estimate the agreement. The Bland-Altman plots the differences (WKYMVM-fMLF) in Mcpm values between the two against the averages (peak values of (WKYMVM+fMLF)/2). Line of equality is presented as black dashed line. The 95% upper and lower CI of limits of agreement is plotted as black dotted line for both the upper and lower limits of agreement. The regression line of differences is drawn as black line with red lines show the 95% confidence interval. Each FFA2R allosteric modulators response is color-coded to a unique color. (**C**) The peak superoxide activities were determined is projected in a bar graph as the ratio between WKYMVM (2.5 nM) and fMLF (0.5 nM) for all allosteric modulators tested. Six independent experiments with cells isolated from healthy donors were performed and the results are given as mean+SD. One-way ANOVA followed by Dunnett’s post-hoc test was used to calculate significant difference deviation from Cmp58.

##### Direct correlation between the FFA2R transduced neutrophil responses induced by FPR agonists

We performed a comparison of the responses for each allosteric FFA2R modulator as a ratio to enable a more direct way to determine the similarities/differences between the effects of the allosteric FFA2R modulators on the NADPH-oxidase activity (Fig 7C). The responses induced by low concentrations of fMLF and WKYMVM had no significant ratio difference between the peak values of the WKYMVM and fMLF. Taken together, these data suggests that allosteric modulated FFA2R was activated equally well by fMLF and WKYMVM, implying that similar downstream signals that activated FFA2R were generated by both FPR1 and FPR2.

## Discussion

Allosteric modulators specific for free fatty acid receptor 2 (FFA2R) transferred not only the orthosteric FFA2R agonist propionate to a neutrophil activating ligand, but also the responses induced by agonists for P2Y_2_R (receptor for ATP), PAFR (receptor for PAF) and the FPRs (FPR1, receptor for fMLF; FPR2, receptor for WKYMVM) were positively modulated. All receptors belong to the family of GPCRs. According to prevailing perception, GPCRs are activated when an orthosteric agonist binds to a receptor site exposed on the cell surface, and this binding starts a G protein signaling cascade that is initiated by the structural changes of the cytoplasmic receptor domains. Activation of the allosterically modulated FFA2R may also be achieved by receptor cross-talk/transactivation mechanisms. This mode of activation of FFA2R is initiated down-stream of P2Y_2_R, PAFR and the FPRs, by signals generated by these receptors on the inside of the plasma membrane. We recently reported binding/signaling/activation-data which show that FFA2R has two distinctly different allosteric receptor sites (9-11). Although the basic structural characteristics the two FFA2R sites remain to be determined, a number of allosteric FFA2R modulators have been described, that selectively bind to one or the other of the two allosteric sites (22, 28). Some of these allosteric FFA2R modulators have been used to investigate the link (if any) between activation of FFA2R by the classical out-side-in signaling mechanism and the intracellular receptor cross-talk signaling mechanism. The data presented herein support the hypothesis that the mechanism by which the neutrophil NADPH-oxidase is activated by FFA2R differs when achieved through agonist binding to the orthosteric site, and when achieved by signals generated down-stream of a cross-talking/transactivating neutrophil receptor.

According to the generally accepted model for how an allosteric GPCR modulator works, binding of a positive modulator either affects the orthosteric binding site and transfers the recognizing receptor to a state characterized by an increased affinity/reduced energy barrier for the conformational change (from nonsignaling to signaling) induced by the agonist, and/or a structural change of the signal transducing receptor domains that more easily/potently respond when the receptor is activated (37, 38). Irrespectively of the precise mechanism, allosteric GPCR modulation should by definition solely affect receptor-mediated responses induced by agonists that are recognized by the modulated receptor. However, the data presented suggests this restriction for allosteric modulators is not valid for FFA2R. The activation of the neutrophil NADPH-oxidase is induced by ATP, PAF as well as low nanomolar concentrations of FPR agonists, all depends on an allosterically modulated FFA2R. This is evident from the fact that FFA2R specific antagonists inhibit this receptor cross-talk response and no allosteric receptor cross-talk activation is achieved in cells lacking FFA2R (10, 11, 19). The precise signaling mechanism by which the neutrophil GPCRs (i.e., P2Y_2_R, PAFR, FPR1 and FPR2) activate the allosterically modulated FFA2R has not been disclosed, but the fact that the receptor cross-talk activating signal(s) generated by P2Y_2_R and PAFR require a functional Gα_q_ subunit, suggests that the receptor cross-talk signals are generated down-stream of the G protein and activate FFA2R from the cytosolic side of the plasma membrane. Although the FPRs probably couple to a Gα_i_ containing G protein (35), similar postulations have earlier been made regarding the FPR1 and FPR2 receptor cross-talk transactivation signal(s).

FFA2R has two distinct different binding sites for allosteric modulators and the two modulators Cmp58 and AZ1729 bind each of these sites (9, 11). At the same time as the three binding site model (one orthosteric and two allosteric sites) was confirmed, several new allosteric FFA2R modulators were identified. These modulators were shown to bind either to the Cmp58 or to AZ1729 site, and to affect the FFA2R response induced by propionate to varying degrees. A selection of these modulators was used to determine the correlation between the two different FFA2R activation mechanisms; that is outside-in activation induced by the orthosteric FFA2R agonist and inside-in receptor cross-talk FFA2R activation.

The data presented showed that there was a moderate positive overall correlation between the propionate and ATP induced responses when all allosteric modulators were included. The correlation/agreement was determined from Pearson and Bland-Altman plots, respectively. Generally Bland-Altman plots are used to show the agreement between two activation processes but unfortunately, there are no values for limits of agreement that are universal in indicating whether an agreement is good or bad (36). This means that there is no predefined cut-off value to refer to, which means experimental context is crucial. The lack of a robust correlation between the propionate and ATP responses was apparent when the ratios between the two responses were determined separately for the different allosteric modulators. Whereas the ratios between the propionate and ATP induced responses were around one for the two “parent” allosteric FFA2R modulators (i.e., Cmp58 and AZ1729), the values were significantly lower for the other modulators with AZ8703 and AZ1702 being the lower extremes. These data suggest that the allosterically modulated FFA2R can adopt structures that differ in the ability to transduce the signals induced by an orthosteric agonist and the signals generated down-stream of P2Y_2_R, respectively.

The results obtained when PAF was used to activate neutrophils, show that a receptor cross-talk/transactivation of FFA2R was obtained, with similar characteristics as that induced by ATP. Whereas the ratio between the propionate and PAF induced responses was around one for the two “parent” allosteric FFA2R modulators, the values were significantly lower for the other modulators with AZ8703 and AZ1702 being the extremes also here. These data suggest that the allosterically modulated FFA2R can adopt a conformation that is better suited to transduce the signals induced by an orthosteric agonist and the signals generated down-stream of the PAFR. In contrast to the non-activating P2Y_2_R agonist ATP, PAF is a weak activating agonist also in the absence of an FFA2R modulator, but the fact that the cross-talk activation of FFA2R, induced both by PAF and ATP, was inhibited by a Gα_q_ selective inhibitor, suggests that similar (but not identical) signals are generated down-stream of the PAFR and the P2Y_2_R. Accordingly, the correlation/agreement determined from Pearson and Bland-Altman plots is in line with this similarity. Generally, the PAF induced response is more pronounced than the ATP response (a mean bias in Bland-Altman of 16.4), but the similarity was apparent when the ratios between the two responses were determined separately for the different allosteric modulators. No significant differences in the ratios between the PAF and ATP induced responses were identified, suggesting that similar FFA2R signals are generated by the two receptors.

We have earlier shown that provided that FFA2R is allosterically modulated, a response is induced by FPR specific agonists at concentrations were these are unable to activate neutrophils alone. The FPRs, just as FFA2R, have been suggested to couple to a Gα_i_ containing G protein (for a more detailed discussion see (13)), suggesting that the receptor cross-talk/transactivation signals may be generated down-stream of both Gα_i_ and Gα_q_ containing G proteins. The two FPRs share a high amino acid sequence homology (39) and have also been shown to share similar signaling/activation patterns (34, 40). Although the responses induced by the low concentrations of fMLF (FPR1 agonist) as well as by WKYMVM (FPR2 agonist) were fairly low, the ratios between the responses induced by the two agonists were close to one for all allosteric modulators. These data are in agreement with the signaling similarities between the FPRs and suggest that identical intracellular cross-talk signals were generated by these receptors, signals that activate the allosterically modulated FFA2R.

The receptor sites that recognize the allosteric modulators Cmp58 and AZ1729 have not yet been identified, but based on the activation induced by two interdependent allosteric FFA2R modulators, AZ5994 and AZ7004 have been classified as “Cmp58 like” whereas the other four (AZ0688, AZ0682, AZ8703 and AZ1702) have been classified as “AZ1729 like” (9). Structurally, AZ0688 and AZ0682 are similar to the parent AZ1729 while AZ8703 and AZ1702 belong to a dihydroisoquinoline-2-amine series of allosteric FFA2R modulators (9). According to the three binding sites model, the “Cmp58-like” and the “AZ1729-like” allosteric modulators should induce the same response pattern as that for the respective parent compound. This was, however, not what was found; this is illustrated by the fact that the response induced by ATP in AZ7004 sensitized neutrophils was fairly low compared to the response induced by propionate, whereas no such difference was evident for AZ5994. AZ7004 is the *S*-enantiomer of 4CMTB (also known as AMG7703), an earlier described racemic phenylacetamide, classified as an allosteric FFA2R agonist. Despite the fact that the structure of AZ7004 is very similar to that of AZ5994, the response pattern following sensitization with the latter follow that of the parent compound Cmp58. The same response pattern as that seen with AZ1729 should be obtained with different “AZ1729-like” allosteric modulators. Once again this was not what was found; whereas the response patterns induced by ATP and PAF in AZ0688 and AZ0682 sensitized neutrophils were very similar to that with the parent compound AZ1729, no such similarity was evident for AZ8703 and AZ1702.

The proposed signaling mechanism by which the cross-talking receptors for ATP, PAF, fMLF, and WKYMVM activates allosterically modulated FFA2R (see Fig 1), starts with an activation of a G protein down-stream of P2Y_2_R, PAFPR, FPR1/FPR2 and the signal(s) generated then activates FFA2R from the cytosolic side of the membrane, and the receptor cross-talk/transactivation signals may be generated down-stream of both Gα_i_ and Gα_q_ containing G proteins. These data suggest similar but possibly not identical signals are generated downstream the different G proteins, but the precise nature of these receptor cross-talk/transactivation signals remains to be determined.

Taken together, our data showing that allosteric FFA2R modulators affect not only the activating potential of propionate, the natural (orthosteric) agonist of the modulated receptor, but also that of several other inflammatory mediators, raises question not only about the regulatory roles of FFA2R in inflammatory settings, but also the mechanisms by which neutrophil GPCRs communicate to activate the ROS generating NADPH-oxidase. In addition, the activation characteristics of allosterically modulated FFA2Rs depend not only on the involvement of one or the other the two different allosteric receptor sites, but also on the nature of the allosteric modulator. Accordingly, the model system with FFA2R together with the different transactivating neutrophil receptors, will be useful in future studies of the allosteric modulation phenomenon and to increase our understanding about the biology of GPCRs biology and the potential of these receptors a drug target to regulate inflammation.

## Abbreviations

CATPB: (*S*)-3-(2-(3-chlorophenyl)acetamido)-4-(4-(trifluoromethyl)-phenyl)butanoic acid an FFA2R antagonist/inverse agonist;
CL: chemiluminescence;
Cmp1: (*R*)-3-benzyl-4-(cyclopropyl(4-(2,5-dichlorophenyl)thiazol-2-yl)amino)-4-oxobutanoic acid a small molecule orthosteric FFA2R agonist;
Cmp58: an allosteric FFA2R modulator;
AZ1729: an FFA2R modulator;
FFA2R: free fatty acid receptor 2;
GPCR: G-protein coupled receptor;
HRP: horse radish peroxidase;
ns: no significant difference;
KRG: Krebs-Ringer Glucose phosphate buffer ;
ROS: reactive oxygen species;
TNF-α: tumor necrosis factor α

## Authorship contributions

*Participation in research design*: Lind, Forsman, Dahlgren

*Conducted experiments:* Lind

*Provided experimental tools*: Granberg

*Performed data analysis:* Lind

*Planned the research and wrote the manuscript:* Lind, Dahlgren

*Supervised the research:* Forsman and Dahlgren

